# Constructing phylogenetic relationship based on the independent selection law of genome sequences

**DOI:** 10.1101/2021.03.20.436244

**Authors:** Li Xiaolong, Li Hong, Yang Zhenhua, Zhang Zefeng

## Abstract

Exploring the composition and evolution regularity of genome sequences and constructing phylogenetic relationship by alignment-free method in genome level are high-profile topics. Our previous researches discovered the CG and TA independent selection law s existed in genome sequences by analysis on the spectral features of 8-mer subsets of 920 eukaryote and prokaryote genomes. We found that the evolution state of genomes is determined by the intensity of the two independent selections and the degree of the mutual inhibition between them. In this study, the two independent selection patterns of 22 primate and 28 insect genome sequences were analyzed further. The two complete 8-mer motif sets containing CG or TA dinucleotide and their feature of relative frequency are proposed. We found that the two 8-mer sets and their feature are related directly to sequence evolution of genomes. According to the relative frequency of two 8-mer sets, phylogenetic trees were constructed respectively for the given primate and insect genomes. Through analysis and comparison, we found that our phylogenetic trees are more consistent with the known conclusions. The two kinds of phylogenetic relationships constructed by CG 8-mer set and TA 8-mer set are similar in insect genomes, but the phylogenetic relationship constructed by CG 8-mer set reflect the evolution state of genomes in current age and phylogenetic relationship constructed by TA 8-mer set reflect the evolution state of genomes in a slight earlier period. We thought it is the result that the TA independent selection is repressed by the CG independent selection in the process of genome evolution. Our study provides a theoretical approach to construct more objective evolution relationships in genome level.

## INTRODUCTION

With the explosive growth of the complete genome sequencing data of species, it has become a study focus to discuss the evolutionary relationship of species in genome levels. The genome sequence contains all of the information about the sequence composition and evolution. The occurrence frequency of k-mers in genome sequence is considered to be an ideal feature set to reveal genome information. It has been found that the frequency of k -mers in DNA sequences is non-random. Based on this feature, functional fragments and functional regions in DNA sequences were analyzed and predicted. Chan and Kibler u sed 6-mers to predict cis-regulatory motifs (Chan et al. 2005). Li and Lin used the k-mers (2≤*k*≤9) to predict the promoter region (Li et al. 2006; Lin et al. 2011). Zhang et al. used k-mers to predict the binding site (Zhang et al. 2011) of locust DNA. Hariharan et al. found that different k -mers were closely related to the diversity of functional fragments (Hariharan et al. 2013). Guo et al. used k-mers to study the nucleosome localization (Guo et al. 2014). Some other studies mainly focused on probabilistic models of k-mers distribution, rare k-mers and rich k-mers fragments and their distribution in chromosomes (Csűrös et al. 2007; Tuller et al. 2007; Hao et al. 2000; Subirana et al. 2010 ; Hampikian et al. 2007; Hariharan et al. 2013; Yu et al. 2013; Chae et al. 2013; Yang et al. 2012; Chikhi et al. 2014), the good results have been obtained in searching for some gene regulatory fragments from k -mers (D’haeseleer at al. 2006; Bina et al. 2009; Bina et al.2004; Xie et al. 2005; Zhang et al. 2011;).

Some studies have tried to study phylogenetic relationship by the non-random features of k-mer usage to overcome the theoretical defects encountered in constructing phylogenetic relationships by using sequence alignment method. Based on the sequence alignment method to construct phylogenetic relationship generally has gone through three stages. At the earliest, phylogenetic relationships were constructed on the basis of single DNA/RNA or amino acid sequence. Such as cytochrome C protein sequence (Pace et al. 2012), small subunit ribosomal RNA (SSU rRNA) sequence (Woese et al. 1977), elongation factor Tu (EF-TU) (Kamla et al. 1996), heat-shock protein (HSP60) gene (Kwok et al. 1999), the largest subunit of the RNA polymerase II (RBP1) (Hirt et al. 1999), the aminoacyl-tRNA synthetases (AARSs) (Woese et al. 2000), etc. However, the inconsistence problem of these phylogenetic trees cannot be solved due to the difference of the selected functional sequences. In addition, it is not appropriate apparently to use the information of a single segment to represent the information of whole genome sequence. Later, people tried to construct phylogenetic relationship by using a set of functional sequences. For instance, common gene set (Snel et al.1999; Huynen et al. 1999), conserved cluster of homologous genes (Wolf et al. 2001), all protein coding sequences or all protein sequences (Qi et al. 2004; Wei et al. 2004; Qi et al. 2004). Although this method improves the reliability of the phylogenetic relationship, it still cannot resolve the inconsistency of the phylogenetic trees because there is no uniform standard to select a consistent sequence set and to determine the number of these sequences, some functional sequences are not possessed by all species. In addition, the selected sequence set still cannot represent the information of whole genome sequence. Some people have attempted to construct phylogenetic relationship by the alignment of whole genome sequence, but this method is only applicable to mitochondrial genome sequence or very small genome sequence (Li et al. 2001; Li et al. 2002), and it is currently not feasible for larger genomes.

A growing number of attentions has been paid to the information of k-mer frequency in whole genome sequence to construct phylogenetic relationship. Earlier, Nussinov explored the compositional heterogeneity of prokaryotic and eukaryotic DNA sequences based on the 2-mer frequency (Nussinov et al. 1984). Karlin et al. analyzed and compared the compositional differences in phage, bacterial and some eukaryote genome sequences based on the k-mer (*k=* 2, 3, 4) frequency (Karlin et al. 1994 ; Gentles et al. 2001). Chapus et al. proposed the idea of constructing phylogenetic relationship based on whole genome sequences using k-mers frequency difference analysis, and discussed the optimum range of *k* value (Chapus et al. 2005), and Qi et al. used the k-mer features set of genome sequences to construct the phylogenetic relationship in prokaryotes (Qi et al. 2004). There are still some insurmountable problems in th is method. If the total k-mers are selected, the phylogenetic trees are very poor, especially in eukaryotes. If the k-mer set is filtered or screened, it is difficult to determine the definite number of k -mers and the arbitrariness problem in the selection of k-mer number cannot be solved. How to determine the number and the feature of k-mers associated with genome evolution is the biggest challenge encountered currently.

A number of studies have focused on the k-mer spectra of genome sequences and the correlations between k-mer spectrum and genome evolution. K-mer spectrum is the specific label of a genome sequence. Chen et al. found that the 6-mer spectra are different for different genome sequences (Chen et al. 2005). Subsequently, Benny Chor et al. found that the k-mer spectra of prokaryotes, fungi and some lower animals were single -peak distributions, while the k-mer spectra of higher animals such as tetrapod mammals showed a triple-peaked distribution, and speculated that the divergence of k-mer spectra is caused by the interaction between CpG inhibition and G+C content constraint (Chor et al. 2009). In our previous works, we studied the spectrum features of 8-mer subsets in 920 species genomes from prokaryote to eukaryote respectively, and found that the spectra distributions of CG and TA 8-mer subsets are independent single-peaked distribution, while this phenomenon was not found in other XY 8-mer subsets spectra. Through analyzing the spectrum features of these 8-mer subsets, we found that there are two independent selection patterns in genome sequences, defined as the CG and TA independent selection laws. In addition, we found that CG1/CG2 subset motifs are directed evolution, they have specific biological functions (Bao et al. 2012; Zhou et al. 2015). Moreover, the spectral features of CG and TA 8-mer spectra are closely related to genome evolution. Therefore, we’re going to search for a set of k-mers and their features that characterize the sequence evolution of genomes based on the independent selection laws of genome sequences, and provide a more reasonable method to construct phylogenetic relationship. We will verify the reliability of our theoretical method based on the whole genome sequences of 22 primates and 28 insects.

## RESULTS

The independent selection laws and the evolutionary mechanism of genome sequences are the theoretical basis of thi s study (see **Method**). Previous studies have pointed out that the four properties of CG and TA independent selection laws reflect the compositional and evolutionary patterns of genome sequence, and the selection pressure of genome evolution is mainly reflected in CG2/CG1 and TA2/TA1 8 -mers. Based on the 22 primate and 28 invertebrate genome sequences (**Table 1 and Supplemental Table 1**), we evaluated the contribution of CG and TA 8 -mers in the sequence evolution of genomes and explored the 8-mer sets and their feature that related directly to genome evolution, and then, proposed a reliable theoretical method to construct phylogenetic relationship.

**Table 1.**
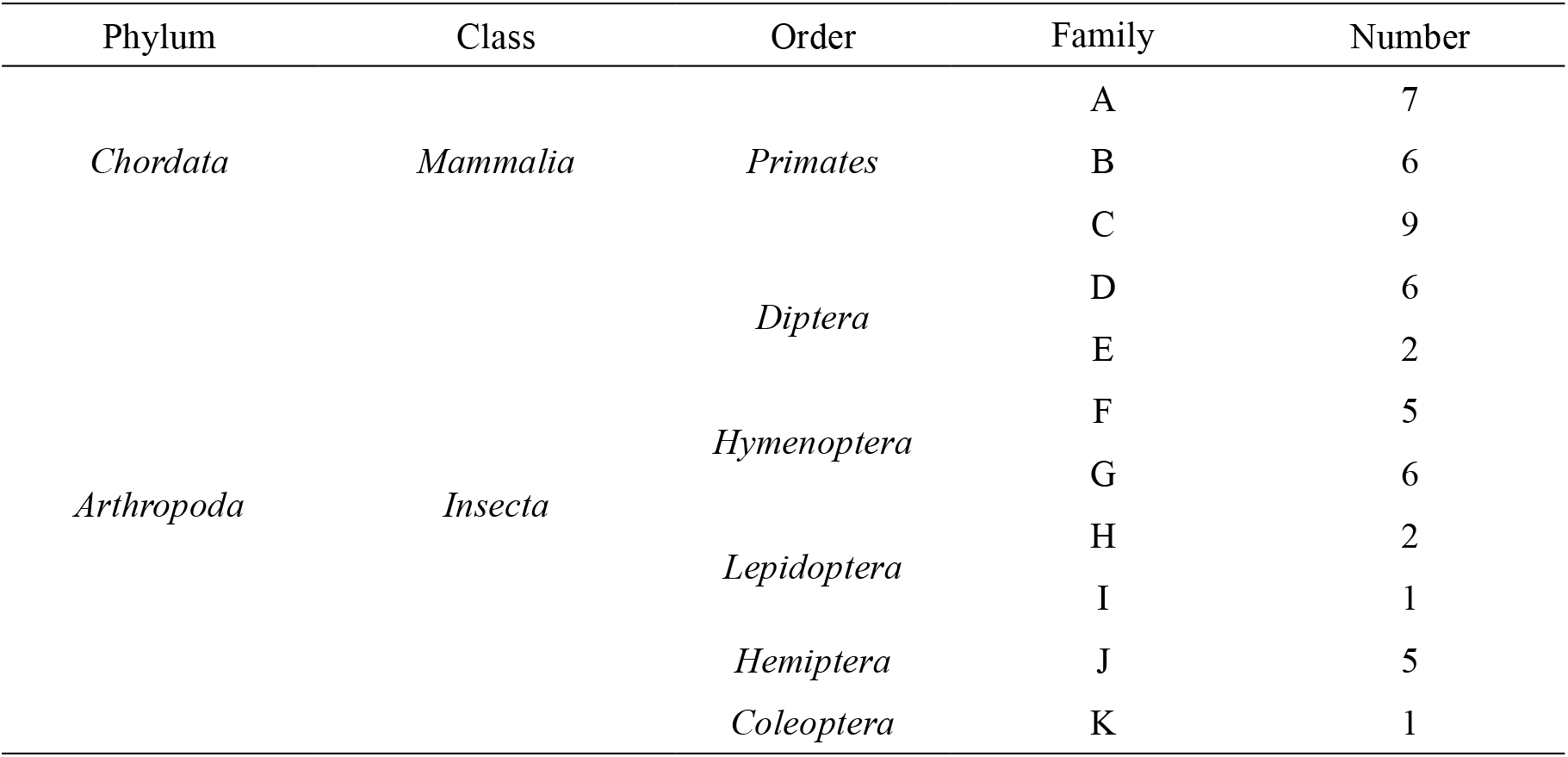
Species classification and genome number

### Intensity distributions of separability and conservatism

According to the characterization method of 8 -mer spectra, the values of separability *δ*_*i*_ and the conservatism *ρ*_*i*_ were calculated for three CG 8 -mer subsets and three TA 8-mer subsets of genome sequences, results are shown in **Supplemental Table 2**. Independent selection law points out that there is a positive correlation between separability and conservatism. Therefore, only the distributions of separability *δ*_*i*_ are presented in **Fig. 1(A, B, C)**.

**Figure 1.**
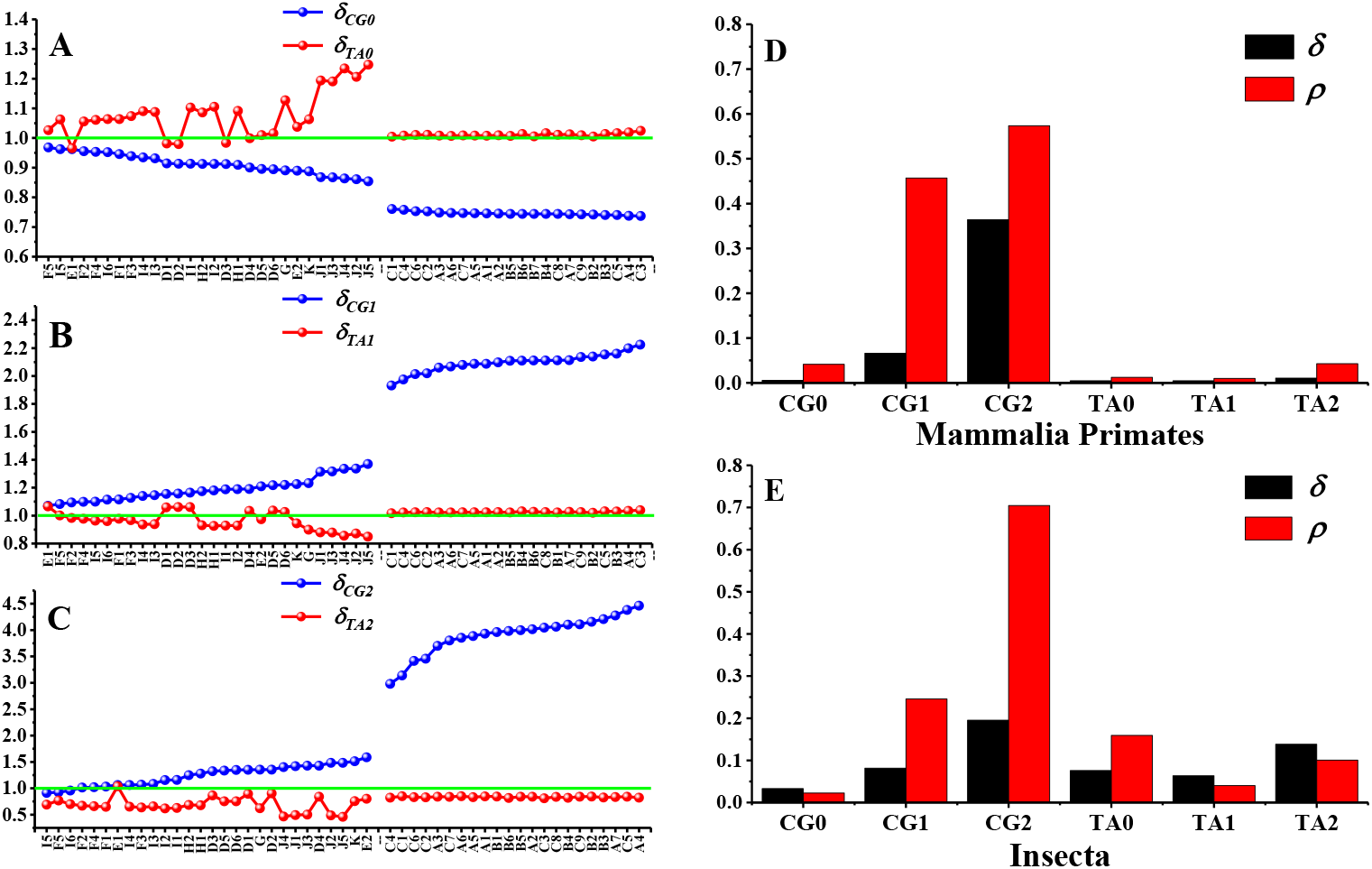
The separability distributions (left) and the standard deviations of the separability and conservatism values in primate and insect genome groups (right). (**A**) The separability distributions of CG0 and TA0 8-mer spectra. (**B**) The separability distributions of CG1 and TA1 8 -mer spectra. (**C**) The separability distributions of CG2 and TA2 8 -mer spectra. (**D**) The standard deviations of the separability and conservatism values for three CG and three TA 8 -mer spectra in primate group. (**E**) The standard deviations of the separability and conservatism values for three CG and three TA 8 -mer spectra in insect group.

By analysis separability and conservatism values in **Fig. 1(A, B, C)** and **Supplemental Table 2**, we found that separability and conservatism values of CG1 and CG2 8-mer spectra is generally higher than that of TA1 and TA2 8-mer spectra in insect genomes, the mutual inhibition relationship does exist for the two parameters between CG1 and TA1 8-mer subsets and between CG2 and TA2 8-mer subsets. In primates, separability and conservatism values of CG1 and CG2 8-mer spectra are higher obviously than that in insect. The two parameters of TA1 and TA2 8-mer spectra are nearby the value *δ*_*i*_=1, which indicates that the phenomenon of TA independent selection has disappeared in primate genomes. The result validated further our previous conclusion (Yang et al. 2020).

We found that the distributions of separability and conservatism values of CG0 or TA0 8-mer spectra are exactly opposite to that of the 8-mer spectra containing CG or TA dinucleotide. That means the distributions of CG0/TA0 8 -mer spectra are also not random. We thought that the opposite distribution features for CG0 and TA0 8-mer spectra are due to the non-random sampling. According to statistical theory, when the 8-mers with low frequency are extracted from the total 8-mer population, the average frequency of the remaining samples must increase, and vice versa. T he distribution features of CG0 and TA0 8-mer subsets is the result of non-random sampling, and that the CG0/TA0 8-mers themselves are not related directly to the evolutionary difference of genome sequences.

In order to investigate the sensitivity of separability and conservatism parameters to genome divergence, the standard deviation distributions of the two parameters for three CG 8-mer subsets and three TA 8-mer subsets were analyzed in primate and insect genomes (**Fig**.

**1D, E**). Actually, the separability reflects the overall frequency variation of the 8 -mers in its subset and the conservatism reflects the preferential feature of 8 -mer usage or the sequential relationship of the 8 -mers in its subset. In combination with the results in **Fig. 1(A, B, C)**, it can be seen that the conservatism parameters of CG1/CG2 8-mer spectra are significantly higher than that of the separability in primate s and in insects, which indicated that the sequence evolution of genomes is reflected not only in the variation of the frequencies of the 8-mers containing CG or TA dinucleotide, but more importantly in the constraints on the sequential relationship of these 8-mers in 8-mer subset.

In a word, the independent selection law reveals the evolution regularity of genome sequences. The features of the 8-mers containing CG or TA dinucleotide correlate directly to the evolution state of genome sequences and the features of CG0 and TA0 8-mers have no direct relationship with sequence evolution of genomes.

### Construction of phylogenetic trees

#### Selection of evolutionary characteristic parameters

Based on the independent selection law of genome sequence, it was found that the 8 -mer motifs containing CG or TA dinucleotide are the core motifs that embody the evolution state of genome sequence, while CG0 and TA0 8-mers are not related directly to genome evolution. Therefore, we selected the 8-mer sets containing CG dinucleotide and containing TA dinucleotide as two kinds of motif sets to characterize the evolution state of genome sequences. The number of the selected 8-mer set is 21468+3523=24991. How to determine the feature of the selected 8-mer set is another key problem. Generally, people used the k-mer probability to characterize the phylogenetic relationship, but the results usually were not satisfied. The feature of k-mer actual probability is not the suitable feature in constructing the phylogenetic relationship. Because the spectral distributions of 8 -mer subsets are not the normal distribution, it seems like the Poisson distribution and the frequencies of some 8-mers in its subset are much large. Using the 8 -mers with extremely high frequency or extremely low frequency to stand for the 8 -mers’ contribution to sequence evolution is bound to exaggerate or weaken the contribution of the 8 -mers. We believe that the contribution of each core motif to genome evolution should be equally weighted. Our analysis in above section pointed out that the sequential relationship in terms of the 8 -mer frequency is more important than the 8-mers’ actual frequency in 8 -mer subset. Considering the sequential relationship and the frequency factors, the actual frequency of the 8 -mers in its subset were transformed as the follow method: the 8 -mer is arranged in order in terms of the 8 -mer frequency from small to large in the subset and the sequence number is used to represent the frequency feature of 8-mers, which is called the relative frequency of 8 -mers. The advantage of the feature is highlighted the sequence relationship of the 8 -mer in its subset and partly eliminated the unequal weight status of the 8 -mers in contributing to genome evolution. The relative frequency is independent of the size of genome sequence so that the relative frequency of 8-mers can be compared in different sequences. Finally, the relative frequencies of 24991 8-mers were selected from total 65536 8 -mers. We considered that the 8-mer set and their relative frequencies include the core information of genome evolution.

### Evolutionary relationship of primates

The phylogenetic trees of 22 primate genomes were constructed by the selected feature set. We knew that only CG independent selection mode is existed in primate genomes. In order to verify the reliability of our feature set, 8-mer motifs are divided into different sets: (1) Total 8-mers, namely the 8-mers in CG0+CG1+CG2 subsets (*N=N*_0_*+N*_1_*+N*_2_*=* 4^8^=65536). (2) The 8-mers in CG1 subset (*N*_1_=21468). (3) The 8-mers in CG2 subset (*N*_2_=3523). (4) The 8-mers in CG1+CG2 subsets (*N*_1,2_*=N*_1_*+N*_2_=24991). In primate genomes, TA 8-mer sets are not considered here because TA independent selection has disappeared. Then, the 8-mer’s actual frequency in subset is transformed into its relative frequency. According to the method, the distance matrices among different species were established and using Mega7 software to obtain the phylogenetic trees. Finally, the phylogenetic tree s of primates as currently recognized are obtained on NCBI and UCSC websites. And it was used as reference s to evaluate the accuracy of our phylogenetic tree s. Phylogenetic trees of 22 primate genomes were obtained from the four types of 8 -mer subsets, results are shown in **Fig. 2(A, B, C, D)**, and reference phylogenetic tree of primates was shown in **Fig. 2E**. The 22 primate species include three genera: apes, monkeys and proto-monkeys, they are marked by B, A and C in **Fig 2**. We can see that the four phylogenetic trees showed the differences in genera levels. Based on the total 8 -mers, the phylogenetic tree is basically the same as that of the reference phylogenetic tree besides *Macaca larvatus* (A4) (**Fig. 2A**). *Macaca larvatus* is isolated earlier than other A and B species, which is inconsistent w ith the reference phylogenetic tree. The phylogenetic tree gotten by CG2 8-mers has the worst effect (**Fig. 2C**). There are two differences compared with the recognized phylogenetic tree. One is that species C8 and C9 are not clustered in the one branch, they are separated earlier than species A and B, the other is that species C7 is separated at the earliest and not in one branch with other species. The phylogenetic trees gotten by CG1 8-mers and CG1+CG2 8-mers are exactly the same (**Fig. 2B, D**), and they possess the best similarity with the reference phylogenetic tree. In the phylogenetic tree constructed by CG1+CG2 8-mers (**Fig.2 D**), the species A and B are clustered in a main branch, C9 and C8 is in one branch and form a main branch with other species C. In addition, the species C7 and species (C4, C5) are not clustered into a single branch in reference phylogenetic tree, but they are in one branch in the phylogenetic tree gotten by the CG1+CG2 8-mers. Actually, species C7, C4 and C5 should be in one branch. From this point of view, the phylogenetic tree gotten by the CG1+CG2 8-mers has the best quality.

**Figure 2.**
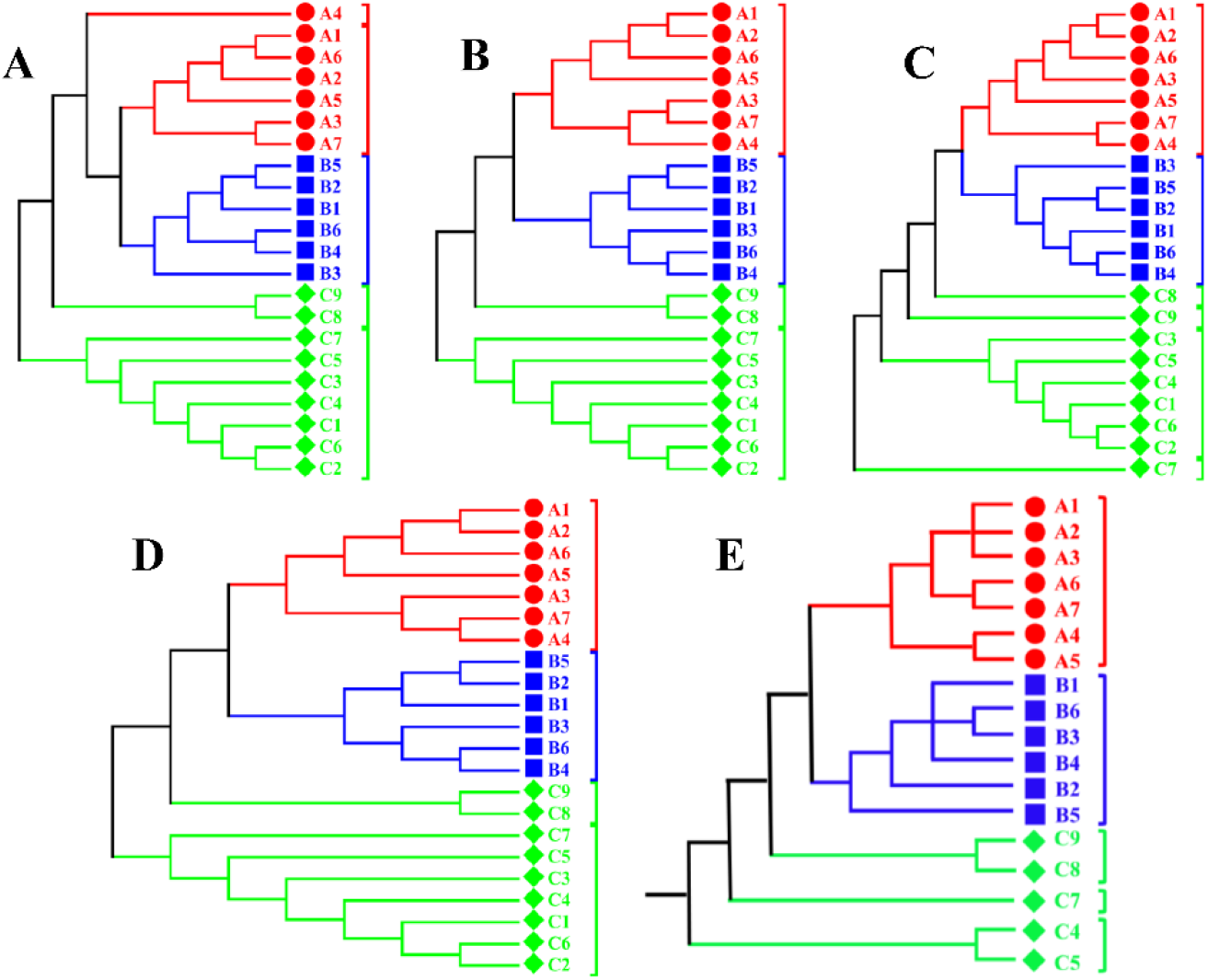
Phylogenetic trees of primates. (**A**) Total 8-mers. (**B**) 8-mers in CG1 subset. (**C**) 8-mers in CG2 subset. (**D**) 8-mers in CG1+CG2 subsets. (**E**) Reference phylogenetic tree.

In our opinion, it is not appropriate to build phylogenetic tree based on the total 8-mers. Independent selection law indicates that CG1/CG2 8-mers are the core motifs that relate directly to the sequence evolution of genomes. Although the 8-mers in CG0 subset can also reflect the information of genome evolution, the information is actually caused by non-random sampling constraints. We thought that there is not new information to be provided by the features of CG0 8 -mers in sequence evolution of genomes. If these motifs are taken into account, it will generate strong background noise, which will inevitably affect the accuracy of the phylogenetic relationship. We considered that the presence of A4 species appeared in the abnormal position (**Fig. 2A**) is caused by the background noise generated by CG0 8-mers.

The phylogenetic tree constructed by CG2 8-mers are defective. The reason is that the number of CG2 8-mers (3523) as the core motifs is extremely small, accounting for only 14% of the core motifs or sensitive motifs (CG1+CG2 subsets). A phylogenetic tree derived from a small number of sensitive motifs is certainly incomplete. The CG1 8-mers account for 86% of sensitive motifs, therefore, it is self -evident that the corresponding phylogenetic tree has high consistency with that by CG1+CG2 8-mers. The above results indicate that using the relative frequency of CG1+CG2 8-mers as the feature parameters to characterize genome evolution is the best choice.

### Evolutionary relationship of insects

We knew that there are both the CG independent selection and the TA independent selection patterns in insect genomes. Based on the conclusion in primates, we are no longer to consider the effect of CG2 and TA2 8-mers. The 8-mer motifs are divided into only three sets:

(1) Total 8-mer motifs. (2) The 8-mers in CG1+CG2 subset. (3) The 8-mers in TA1+TA2 subset. The actual 8-mer frequency is converted into the relative frequency according to the method provided, and the distance matrix between different species is given by the definitions in method. The corresponding phylogenetic trees are shown in **Fig. 3**.

**Figure 3.**
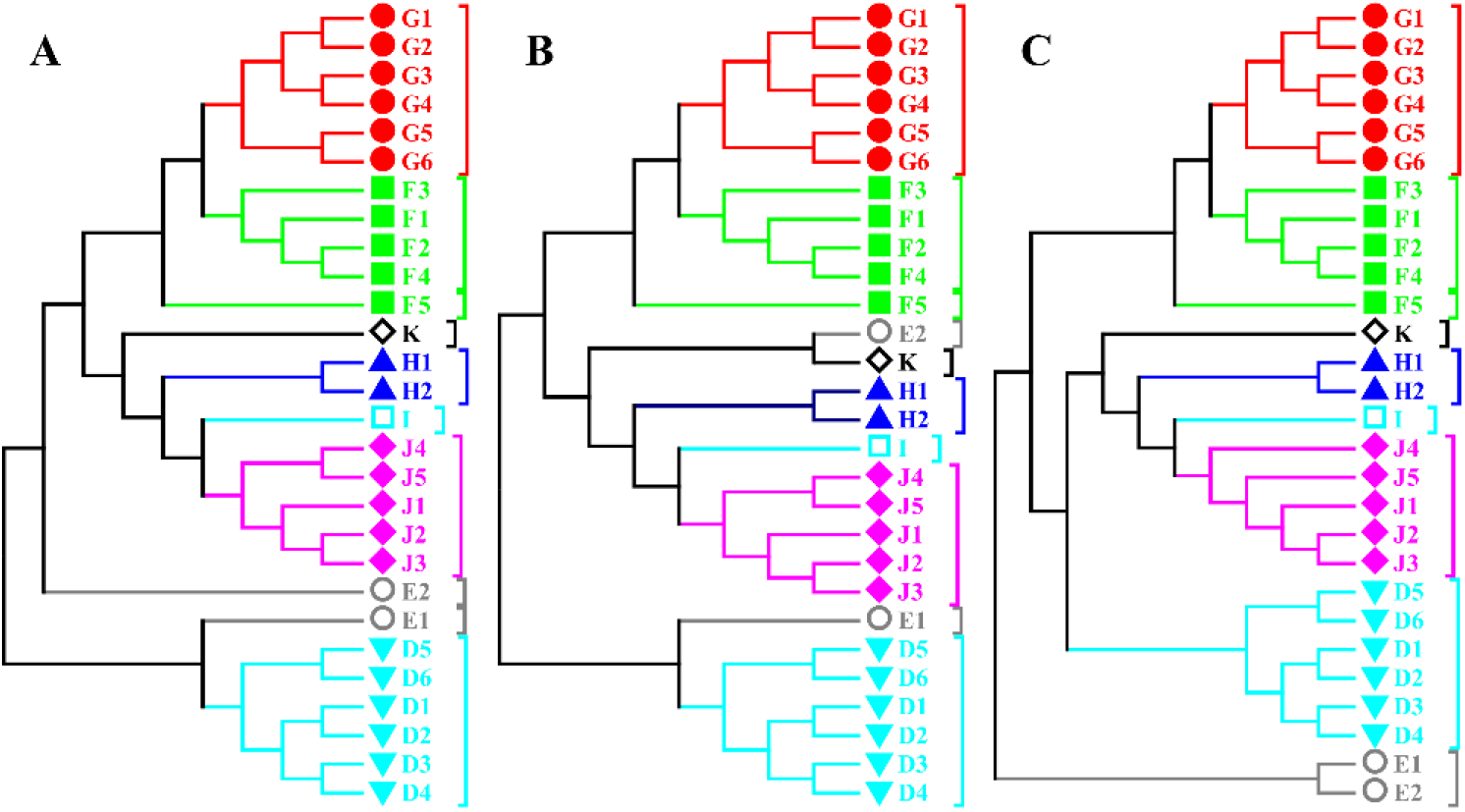
Phylogenetic tree of insecta. (**A**) total 8-mers. (**B**) 8-mers in CG1+CG2 subset. (**C**) 8-mers in TA1+TA2 subset.

Generally, the three phylogenetic trees are almost identical. Species in *Hymenoptera Formicidae* (I), *Hymenoptera Apidae* (F), *Diptera Drosophilidae* (D), *Hemiptera Aphididae* (J) and *Lepidoptera Bombycidae* (H) families are well clustered together. At the order level, the species in family I and family F are clustered in one main branch. The species in family J, H, K and G are clustered in another main branch, and the species in family D is located on the third main branch. The results are agree with the accepted species classification s (Rasnitsyn et al. 2002; Grimaldi et al. 2005). In the three phylogenetic trees, the main differences appeared in the classification of *Contarinia nasturtii* (E2) and *Anopheles gambiae* (E1), which locations are different in the three phylogenetic trees.

The phylogenetic tree constructed by total 8-mers are not substantially different from that by CG1+CG2 8-mers and by TA1+TA2 8-mers (except for E1 and E2). The results reconfirmed again the conclusion that the core motifs for sequence evolution are CG1 +CG2 or TA1+TA2 8-mers and not the CG0/TA0 8-mers.

Except for the E1 and E2 species, the two phylogenetic trees constructed by CG1+CG2 8-mers (**Fig. 3B**) and TA1+TA2 8-mers (**Fig. 3C**) are identical. That is, the two types of 8-mer feature sets are equivalent. We considered that this is a direct manifestation of the mutual inhibition relationship between CG and TA independent selections in insect genomes, and the conclusion is determined by the evolution mechanism of genome sequences.

The two kinds of mosquitoes E1 and E2 are clustered in different positions in the phylogenetic trees constructed by CG1+CG2 8-mers and TA1+TA2 8-mers. We thought that the difference is caused by the difference of the evolution information which includes in the two kinds of 8-mer sets. Our previous study (Yang et al. 2020) shown that the mutual inhibition relationship between CG and TA independent selections originated the evolution pressure of aerobic environment. Early organisms usually took two kinds of independent selection patterns to adapt the aerobic environment. The one is to enhance the intensity of CG independent selection and inhibit the intensity of TA independent selection in the process of genome evolution, such as invertebrates. The other one is to enhance the intensity of TA independent selection and inhibit the intensity of CG independent selection in the process of genome evolution, such as green alga. With the enhancing of genome evolution level s in animals and plants, the intensity of CG independent selection is increasing and of TA independent selection is decreasing. The conclusion means that the aerobic environment is the direct cause to result in the intensity enhancement of CG independent selection. Hence, CG independent selection is the driving force and TA independent selection is changed passively under the mutual inhibition restrains. Thus, there should be a time lag effect between CG and TA independent selections i n terms of reflecting genomic evolutionary information. We considered that the passive response of TA independent selection must reflect the legacy information of the genomes in slightly earlier time.

In the phylogenetic tree constructed by TA1+TA2 8-mers (**Fig. 3C**), the two mosquitoes *Anopheles gambiae* (E1) and *Contarinia nasturtii* (E2) are clustered together to form an individual branch, but they are not clustered in one branch in the phylogenetic tree constructed by CG1+CG2 8-mers (**Fig. 3B**). According to the above conclusion, w e believed that the two kinds of mosquitoes were homologous at a slightly earlier period, and this information is included in TA1+TA2 8 -mer features. Driven by different living environment, the evolution rates of the two kinds of mosquitoes are quite different, resulting in a rapid diverged from each other’s evolution patterns. This information is included in CG1+CG2 8-mer features. The result also reflected the complexity and specificity of the two kinds of mosquitoes in genome evolution. We could conclude that the information of CG1+CG2 8-mers reflects the current evolutionary state of genomes, whereas the information of TA1+TA2 8-mers reflects the evolutionary state of genomes at a slightly earlier period or it can reflect the legacy information of genomes at earlier period.

## CONCLUSION AND DISCUSSION

Based on the CG and TA independent selection laws of genome sequences, which found by our previous studies, we analyzed the frequency spectral features of various 8-mer subsets in 22 primate and 28 insect genome sequences. The results show that there is CG and TA independent selection patterns in insect genome sequences and the two independent selections have mutual inhibition relationship, but only CG independent selection pattern in primate genome sequences, and TA independent selection pattern has disappeared formally. By analyzing the separability and the conservatism of the spectra in CG and TA 8-mer subsets, we found that the 8-mers containing CG and TA dinucleotide correlate directly to sequence evolution of genomes, while the CG0 and TA0 8 -mers are not related directly to sequence evolution of genomes. For the 8-mer spectra containing CG dinucleotide, the feature of conservatism is more sensitive than that of separability in reflecting the genome divergences. It shows that the key feature is the preferential feature of 8 -mer usage or the sequential relationship of the 8 -mers in its subset, but not the 8-mer frequency itself. So, the relative frequency was proposed. The feature of relative frequency reflects not only the magnitude of 8-mer frequency in some extent, but more importantly it reflects the sequential relationship of the 8-mers in its subset. Thereout, we proposed two sets of parameters to characterize the phylogenetic relationship. The first set of parameters is the 24991 relative frequencies of CG1+CG2 8-mers, and the second set of parameters is the 24991 relative frequencies of TA1+TA2 8-mers. The phylogenetic tree of primates constructed by the relative frequency set of CG1+CG2 8-mers is consistent with the known species phylogenetic relationship, and its reliability is even better than the known phylogenetic relationship. Because CG and TA independent selection patterns exist together in insect species, the two phylogenetic trees constructed by the relative frequency set of both TA1+TA2 8-mers and CG1+CG2 8-mers are equivalent and they are consistent with the known phylogenetic relationship. According to the mutual inhibiti on relationship between CG and TA independent selection s in the process of genome evolution, we speculated that the feature of TA1+TA2 8-mers includes the evolution information of genomes at a slightly earlier period and the feature of CG1+CG2 8-mers reflects the evolution information of genomes at present state.

We proposed a theoretical method in constructing phylogenetic relationships based on the evolution mechanism of genome sequences. Our method has the following features: (1) The definite motif sets are given. In the case of 8 -mer motifs, two sets of 24991 8-mers are determined which correlates directly to sequence evolution of genomes, they are the 8-mers containing CG or TA dinucleotide. Our method solves the puzzled problem of arbitrariness in k-mer screening and eliminates the influence of noise generated by the 8-mers that are not related directly to genome evolution. (2) The more objective feature of the selected 8-mers is proposed. The information of the 8-mer relative frequency includes the characteristics both the separability and the conservatism of the 8-mer spectra. The feature of the relative frequency highlights the character of equal right for the 8-mers with very high and very low actual frequencies to contribute to sequence evolution of genomes. (3) The two kinds of selected 8-mer sets should reflect different evolutionary states in different period. We can use the two kinds of the 8-mer sets to assess more clearly the evolution relationships of genomes. Certainly, the conclusion needs to be validated by more examples.

To construct more objective phylogenetic relationship s, we validated the reliability of our theoretical method only in primate and insect genomes. Our previous study indicated that, besides vertebrate genomes, both the CG and TA independent selections exist in the genomes that include plant, fungus, bacteria and archaea and the two independent selections have mutual inhibition relationship. In vertebrate, only CG independent selection is obvious and TA independent selection is disappearing with the evolution level increasing of genomes. So, we considered that our method has the species universality. Our method has not any subjective constraint on genome sequences, the selected motif sets and the motif feature are uniform in constructing phylogenetic relationships. We will further validate the reliability of our theoretical method in all species from eukaryotes to prokaryotes. At the same time, we hope relevant researchers to participate in this validation work and finally set up a standardized and universal methods in constructing phylogenetic relationships.

## DATASET AND METHOD

### Dataset

The complete genome sequences and annotation information of 22 primate species in mammalia and 28 invertebrate species in insecta were obtained from NCBI (National Center for Biotechnology Information) (https://www.ncbi.nlm.nih.gov/). The sex chromosomes were not included in our dataset. The species information was listed in **Table 1** and the detailed information was listed in **Supplemental Table 1**.

## Method

### Selection of *k* value in k-mer

In k-mer spectra analysis, we chose *k=* 8. This choice is based on the considerations. Firstly, when *k*>6, statistical results show that the distributions of k-mer spectra of genome sequences tends to be stable. Secondly, the statistically significan ce should be ensured for the frequency of k-mers occurrence in genome sequence. If *k* value is too large, the number of k-mer motifs (*N=* 4^*k*^) is too much, and many k-mers appear with zero frequency, which will result in loss of information. Beny Chor (Chor et al. 2009) proposed a formula for minimum *k* value estimation. It is *k =* 0.7*log*_4_ *L, L* is the length of DNA sequence. After examination, *k* value is greater than 8 for eukaryotic genomes and greater than 6 for prokaryotic genomes. Accordingly, we chose *k=* 8 in our study.

### 8-mer spectrum

For a given genome sequence, the frequency of each 8 -mer is obtained by using 8 bp as the window and 1 bp as the step size. If the number of 8 -mers that occur *i* times is *N*_*i*_, the 8-mer relative motif number (*RMN*) is defined as:

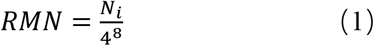

The distribution of the *RMN* value with the 8-mer frequency is called 8 -mer spectrum of the genome sequence.

### Classification method of 8-mers

After obtaining the frequency of each 8-mer in a genome sequence, we classified the 8-mer set into different subsets according to the composition features of 8-mers. The 8-mers containing zero XY (X, Y=A, T, C, G) dinucleotide is classified into XY0 subset, containing one XY dinucleotide is classified into XY1 subset and containing two or more than two XY dinucleotides is classified into XY2 subset. Theoretically, there are 4^8^=65536 8-mer motifs. When X≠Y, there are 40545 8-mers in XY0 subset, 21468 8 -mer in XY1 subset, and 3523 8-mer in XY2 subset. When X=Y, the number of 8 -mers in the three subsets is 44631, 14931 and 5974, respectively. The method is called XY dinucleotide classification method. Thus, total 8-mers are divided into 16 XY classes. In each XY class, there are three XY 8-mer subsets, called XY0, XY1 and XY2. Finally, we could obtain the 8-mer spectra of 48 XY subsets in a given genome sequence.

### Separability and conservatism of 8-mer subset spectrum

For a given 8-mer spectrum, the average and standard deviation of the spectrum distribution are used to characterize the distribution features. In order to compare the distribution position and distribution conservativeness among different 8-mer subset spectra, the separability and conservatism values are defined.

(1) Separability

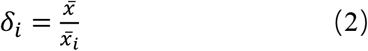

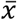 is the average frequency of total 8-mer spectrum, called random center.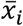 is the average frequency of the *i*-th 8-mer subset spectrum. δ_*i*_ represents the separability of the distribution position of the *i*-th 8-mer subset spectrum relative to that of total 8-mer spectrum. If δ_*i*_>1, it indicates that the 8-mer subset spectrum is located at the low frequency end of total 8-mer spectrum. If δ_*i*_ = 1, it indicates that location of the 8 -mer subset spectrum does not deviate.

(2)Conservatism

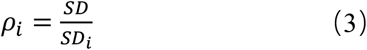

SD is the standard deviation of total 8 -mer spectrum, S*D*_*i*_ is the standard deviation of the *i*-th 8-mer subset spectrum. *ρ*_*i*_ represents the conservatism of the standard deviation of the *i*-th 8-mer subset spectrum relative to that of total 8-mer spectrum. *ρ*_*i*_ > 1, indicating that the conservatism of the 8-mer subset spectrum is higher than that of the total 8-mer spectrum, that means the frequency distribution of the 8-mer subset spectrum is relatively more concentrated.

In the definition of separability and conservatism, the effect of genome size and absolute position of the subset spectrum are eliminated, the two parameters can compare not only the feature difference of different 8-mer subset spectra within a genome, but also the feature difference of 8 -mer subset spectra between different genomes.

### Independent selection law of genome sequences

Previous work analyzed the intrinsic regularity of the 8-mer subset spectra in 920 genome sequences from eukaryotes to prokaryotes. We found that there is the generalized law in genome sequences, called CG and TA independent selection law s (Yang et al. 2020). The two independent selection laws have four properties:(1) Evolutionary Independence. Only the 8-mer spectra of CG0, CG1, CG2 subsets and TA0, TA1, TA2 subsets form the independent single-peak distribution, while the 8 -mer spectra of other XY0, XY1 and XY2 subsets do not show this phenomenon (**Fig. 4**). (2) Evolutionary Selectivity. The 8-mers in CG1/CG2 and TA1/TA2 subsets evolve in a directed way, while the 8-mers in CG0/TA0 subsets evolve randomly. (3) Evolutionary Correlation. There is a positive correlation between the separability and the conservatism of the 8-mer subset spectra containing CG and TA dinucleotide. (4) Evolutionary Homoplasy. The directed evolving ways are same for the 8-mers in CG1 and CG2 subsets as well as that in TA1 and TA2 subsets. Our study shows that the separability and the conservatism of the 8 -mer subset spectra containing CG and TA dinucleotide reflect the evolutionary state of a genome. While the separability or the conservatism parameters are used to characterize the intensity of independent selections, results show that there is a mutual inhibition relationship between CG and TA independent selections. Thereby, we proposed an evolutionary mechanism of genome sequences: the intensities of CG and TA independent selection s and the mutual inhibition relationship between the two independent selections determine the evolutionary state of a genome sequence. We found that the intensity of the CG independent selection is positively correlated and the TA independent selection is negatively correlated with the evolution level s of animal and plant genomes.

**Figure 4.**
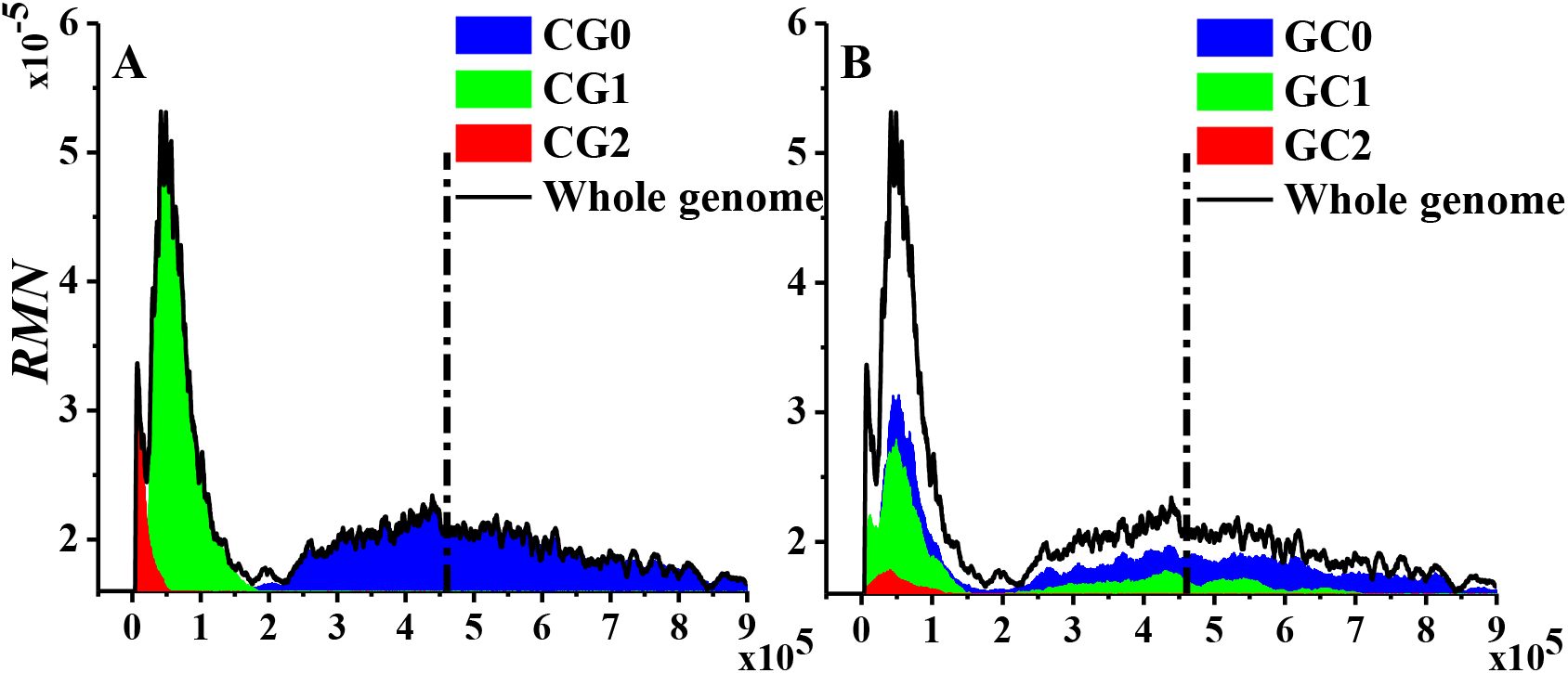
8-mer spectra of *Pan troglodytes* genome sequence. The vertical bar is the average frequency of total 8-mers. (**A**) Spectra of three CG 8-mer subsets and total 8-mers. (**B**) Spectra of three GC 8-mer subsets and total 8-mers.

### Construction of phylogenetic trees

In order to construct the evolutionary relationship, the feature difference of selected 8-mers were used to construct the distance between two species genomes. The definition is as follows:

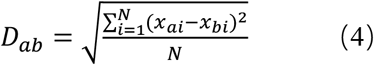

Where *a* and *b* represent two genomes, *x*_*ai*_ is the *i*-th 8-mer feature in genome *a*, and *x*_*bi*_ is the *i*-th 8-mer feature in genome *b. N* is the total number of the selected 8-mers.

Matrix element *D*_*ab*_ is used to construct the distance matrix for a given genome group. Using Mega7 software (http://www.megasoftware.net), the phylogenetic tree is constructed by Neighbor-Joining method.

## Acknowledgements

This work was supported by the National Natural Science Foundation of China (No. 31860304).

These authors contributed equally: Li Hong and Li Xiaolong.

L. H. and L. X.L. conceived the study, performed the data analysis and wrote the manuscript.

Y. Z.H. and Z. Z.F. investigated the optimum characteristics of sets.

## References

Chan BY, Kibler D. 2005. Using hexamers to predict cis-regulatory motifs in Drosophila. BMC Bioinformatics 6(1): 262.

Li QZ, Lin H. 2006. The recognition and prediction of sigma70 promoters in Escherichia coli K-12. Journal of Theoretical Biology 242(1): 135–141.

Lin H, Li QZ. 2011. Eukaryotic and prokaryotic promoter prediction using hybrid approach. Theory in Biosciences 130(2): 91–100.

Zhang Y, Wang XH, Kang L. 2011. A k-mer scheme to predict piRNAs and characterize locust piRNAs. Bioinformatics 27(6): 771–776.

Hariharan R, Simon R, Pillai MR, Taylor TD. 2013. Comparative Analysis of DNA Word Abundances in Four Yeast Genomes Using a Novel Statistical Background Model. PLoS One 8(3): e58038.

Guo SH, Deng EZ, Xu LQ, Ding H, Lin H, Chen W, Chou KC. 2014. iNuc-PseKNC: a sequence-based predictor for predicting nucleosome positioning in genomes with pseudo k -tuple nucleotide composition. Bioinformatics 30(11): 1522–1529.

Csűrös M, Noé L, Kucherov G. 2007. Reconsidering the significance of genomic word frequencies. Trends in Genetics 23(11): 543–546.

Tuller T, Chor B, Nelson N. 2007. Forbidden penta-peptides. Protein Sci 16(10): 2251–2259.

Hao BL, Lee HC, Zhang SY. 2000. Fractals related to long DNA sequences and complete genomes. Chaos, Solitons & Fractals 11(6): 825–836.

Subirana JA, Xavier M. 2010. The most frequent short sequences in non-coding DNA. Nucleic Acids Research 38(4): 1172–1181.

Hampikian G, Andersen T. 2007. Absent Sequences: Nullomers and Primes. Pacific Symposium on Biocomputing 12: 355–366.

Hariharan R, Simon R, Pillai MR, Taylor TD. 2013. Comparative analysis of DNA word abundances in four yeast genomes using a novel statistical background mode. Plos One 8(3): e58038.

Yu HJ. 2013. Segmented K-mer and its application on similarity analysis of mitochondrial genome sequences. Gene 518(2): 419–424.

Chae H, Park J, Lee SW, P. Nephew K, Kim S. 2013. Comparative analysis using k-mer and k-flank patterns provides evidence for CpG island sequence evolution in mammalian genomes. Nucleic Acids Research 41(9): 4783–4791.

Yang Y, Nenneth K, Kim S. 2012. A novel k-mer mixture logistic regression for methylation susceptibility modeling of CpG dinucleotides in human gene promoters. BMC Bioinformatics 13(3): S15.

Chikhi R, Medvedev P. 2014. Informed and automated k-mer size selection for genome assembly. Bioinformatics 30(1): 31–37.

D’haeseleer P. 2006. What are DNA sequence motifs? Nature Biotechnology 24(4):423–425.

Bina M, Wyss PJ, Lazarus SA, Shah SR, Ren W, Szpankowski W, Crawford GE, Park SP, Song XC. 2009. Discovering sequences with potential regulatory characteristics. Genomics 93(4): 314–322.

Bina M, Wyss PJ, Ren W, Szpankowski W, Thomas E, Randhawa R, Reddy S, John PM, Pares-Matos EI, Stein A, et al. 2004. Exploring the characteristics of sequence elements in proximal promoters of human genes. Genomics 84(6): 929–940.

Xie X, Lu J, Kulbokas EJ, Golub TR, Mootha V, Lindblad-Toh K, Lander ES, Kellis M. 2005. Systematic discovery of regulatory motifs in human promoters and 3’UTRs by comparison of several mammals. Nature 434(7031): 338–345.

Pace NR, Sapp J, Goldenfeld N. 2012. Phylogeny and beyond: Scientific, historical, and conceptual significance of the first tree of life. PNAS 109(4): 1011–1018.

Woese CR, Fox GE. 1977. Phylogenetic structure of the prokaryotic domain: the primary kingdoms. PNAS 74(11): 5088–5090.

Kamla V, Henrich B, Hadding U. 1996. Phylogeny based on elongation factor Tu reflects the phenotypic features of mycoplasmas better than that based on 16S rRNA. Gene 171(1): 83–87.

Kwok AYC, Su SC, Reynolds RP, Bay SJ, Av-Gay Y, Dovichi NJ, Chow AW. 1999. Species identification and phylogenetic relationships based on partial HSP60 gene sequences within the genus Staphylococcus. International Journal of Systematic Bacteriology 49(3): 1181–1192.

Hirt RP, Logsdon JM, Healy B, Dorey MW, Doolittle WF, Embley TM. 1999. Microsporidia are related to Fungi: evidence from the largest subunit of RNA polymerase II and other proteins. PNAS 96(2): 580–585.

Woese CR, Olsen GJ, Ibba M, Söll D. 2000. Aminoacyl-tRNA synthetases, the genetic code, and the evolutionary process. Microbiology and Molecular Biology Reviews 64(1): 202–236.

Snel B, Bork P, Huynen MA. 1999. Genome phylogeny based on gene content. Nature Genetics 21(1): 108–110.

Huynen M, Snel B, Bork P. 1999. Lateral Gene Transfer, Genome Surveys, and the Phylogeny of Prokaryotes. Journal of Early Adolescence 17(2): 109–128.

Wolf YI, Rogozin IB, Grishin NV, Tatusov RL, Koonin EV. 2001. Genome trees constructed using five different approaches suggest new major bacterial clades. BMC Evolutionary Biology 1(1): 1–22.

Qi J, Wang B, Hao BL. 2004. Whole Proteome Prokaryote Phylogeny Without Sequence Alignment: A K -String Composition Approach. Journal of Molecular Evolution 58(1): 1–11.

Wei HB, Qi J, Hao BL. 2004. Prokaryote phylogeny based on ribosomal proteins and aminoacyl tRNA synthetases by using the compositional distance approach. Science in China (Series C) 47: 313–321.

Qi J, Luo H, Hao BL. 2004. CVTree: a phylogenetic tree reconstruction tool based on whole genomes. Nucleic Acids Research 32(Web Server issue) : 45–47.

Li M, Badger JH, Chen X, Kwong X, Kearney P, Zhang H. 2001. An information-based sequence distance and its application to whole mitochondrial genome phylogeny. Bioinformatics 17(2): 149–154.

Li W, Fang W, Ling L, Wang J, Xuan Z, Chen R. 2002. Phylogeny Based on Whole Genome as inferred from Complete Information Set Analysis. Journal of Biological Physics 28(3): 439–447.

Nussinov R. 1984. Doublet frequencies in evolutionary distinct groups. Nucleic Acids Research 12(3): 1749–1763.

Karlin S, Cardon LR. 1994. Computational DNA Sequence Analysis. Annual Review of Microbiology 48(1): 619–654.

Gentles AJ, Karlin S. 2001. Genome-Scale Compositional Comparisons in Eukaryotes. Genome Research 11(4): 540–546.

Chapus C, Dufraigne C, Edwards S, Giron A, Fertil B, Deschavanne P. 2005. Exploration of phylogenetic data using a global sequence analysis method. BMC Evolutionary Biology 5: 63.

Chen YH, Nyeo SL, Yeh CY. 2005. Model for the distributions of k -mers in DNA sequences. Physical Review E 72(1 Pt 1): 011908.

Chor B, Horn D, Goldman N, Levy Y, Massingham T. 2009. Genomic DNA k-mer spectra: models and modalities. Genome Biology 10(10): R108.

Bao T, Li H, Zhao XQ, Liu GJ. 2012. Predicting nucleosome binding motif set and analyzing their distributions around functional sites of human genes. Chromosome Research 20(6): 685–698.

Zhou DL, Li H, Yang XX. 2015. Distributions of 8 -mer Frequency of Appearence and the Evolution Diversity of 8 -mer Usage in DNA Sequences of Human Chromosome 1. Acta Biophysica Sinica 31(1): 63–64.

Yang ZH, Li H, Jia Y, Zheng Yan, Meng H, Bao T, Li XL, Luo LF. 2020. Intrinsic laws of k -mer spectra of genome sequences and evolution mechanism of genomes. BMC Evolutionary Biology 20(1). doi: 10.1186/s12862-020-01723-3

Rasnitsyn AP, Quicke DLJ. 2002. History of Insects. Kluwer Academic Publishers. Dordrecht, The Netherlands.

Grimaldi D, Engel MS. 2005. Evolution of the Insects. Cambridge University Press. New York.

